# The structure and diversity of the bacterial community in the overlying water of the Yangzong Lake in Yunnan

**DOI:** 10.1101/2024.03.28.587311

**Authors:** Kai Huang, Likun Li, Jin Li, Han Chen, Zhiqiang Ma, Wenlong Ye, Deping Kong, Jun Zhang

## Abstract

Bacteria are critical components of lake ecosystems, and understanding the characteristics of bacterial community structure and diversity indices is of significant importance for the analysis and management of lake ecology. In this study, overlying water samples were collected from Yangzong Lake in May, June, and July 2021. High-throughput sequencing and statistical analysis were employed to investigate the relationships between physicochemical factors, bacterial community structure, and diversity in the overlying water of Yangzong Lake. The results showed significant differences in oxidation-reduction potential among the sampling points. The order of oxidation-reduction potential was as follows: Yangzong DAHE > Yangzong Lake South > Yangzong Lake Middle > Yangzong Lake North, while other physicochemical factors exhibited minor differences. The number of OTUs and bacterial alpha diversity index in the overlying water of Yangzong Lake Middle were higher than those in Yangzong Lake North, Yangzong Lake South, and Yangzong DAHE. The dominant bacterial phyla in the overlying water were Actinobacteriota, Bacteroidota, and Proteobacteria. Actinobacteriota had the highest relative abundance of 34.63% in Yangzong Lake North, while Bacteroidota had the highest relative abundance of 33.79% in Yangzong Lake South, and Proteobacteria had the highest relative abundance of 27.29% in Yangzong Lake Middle. The dominant genera were hgcI_clade, CL500-29_marine_group, and Flavobacterium. Among them, hgcI_clade had the highest relative abundance of 37.56% in Yangzong Lake North, CL500-29_marine_group had the highest relative abundance of 35.22% in Yangzong Lake South, and Flavobacterium had the highest relative abundance of 42.02% in Yangzong Lake Middle. Correlation analysis revealed significant correlations between Campilobacterota at the phylum level and dissolved oxygen (DO) and total phosphorus (TP). At the genus level, Flavobacterium, Limnohabitans, and Pseudarcicella showed significant correlations with DO and TP.

## 1. introduction

Microorganisms in lakes play a crucial role in the cycling of elemental substances and are essential components of aquatic ecosystems (Wu et al., 2017). Bacteria in lake water serve as both consumers and producers, making them key to understanding the structure and function of lake ecosystems (Uehlinger et al., 1986; Wilken et al., 2018). Changes in environmental factors within lakes can influence the diversity and community structure of waterborne bacteria. Previous studies have found that variations in environmental factors such as dissolved oxygen(DO) and total nitrogen( TN) are major contributors to differences in bacterial community characteristics in lakes (Wang et al., 2023). Exploring the effects of environmental factors on bacterial diversity and community structure in lake water can enhance our understanding of material cycling and energy flow in lake ecosystems (Zhou et al., 2022).

The nine major plateau lakes (>30km^2^) in Yunnan Province are significant ecological and environmental functional areas in southwest China, playing a crucial role in the regional economic and social development of Yunnan Province (Zhao et al., 2023; Ma et al., 2022). Yangzong Lake, as one of the nine major plateau lakes, is an important component of the upstream Pearl River and plays a critical role in biodiversity conservation in Yunnan Province and the establishment of an ecological security barrier in southwest China (Zhang et al., 2022). It not only serves as an essential part of the ecosystem but also bears the responsibility of maintaining the ecological security pattern of the upper Pearl River (Zhang et al., 2022). However, due to increasing human activities, Yangzong Lake’s water quality has been contaminated, especially with the arsenic pollution event in 2008, causing significant losses to drinking water, irrigation, and fisheries and greatly impacting the aquatic ecological security of the Yangzong Lake region (Chen et al., 2015). After the arsenic contamination event, approximately 2,000 tons of ferric chloride were introduced to reduce the arsenic concentration in the water, but the overall changes in arsenic concentration across the entire lake remain unclear (Xu et al., 2022).

This research aims to analyze the content of arsenic, bacterial diversity, and bacterial community structure in the overlying water of Yangzong Lake, to understand the environmental risk of arsenic in the lake and deepen our understanding of the aquatic ecosystem in this lake (Chen et al., 2019). Therefore, this study utilizes high-throughput sequencing technology and employs multivariate statistical analysis to analyze the community characteristics of bacteria in the overlying water of Yangzong Lake and investigate the relationship between bacterial community characteristics and physicochemical factors in the water. The results of this study have important implications for the water environmental quality and ecological conservation of Yangzong Lake, and it also aims to provide valuable references for the protection and management of the water environment in the upstream Pearl River.

## 2 Materials and Methods

### 2.1 Study Site Information

The study site is located in Yangzong Lake Scenic Area, Yiliang County, Kunming City, Yunnan Province, China (24°01’∼24°98’ N, 102°59’∼103°02’ E). Yangzong Lake is a freshwater deep lake on the Yunnan Plateau, belonging to the Nanpan River System in the Pearl River Basin. The average annual temperature is 14.5°C, with an average precipitation of 963.5mm (Yuan et al., 2014). The average water level is 1770.6m, the lake has an average length of 12km north to south and an average width of 2.7km east to west. The lake area is 31.7 km^2^, with a maximum depth of 30m and a storage capacity of 6.04×10^8^m^3^. The watershed area is 192 km^2^ (Bai et al., 2019). The main inflowing rivers are the Yangzong DAHE and Qili River in the south, as well as temporary streams on the east and west sides. The only outflow is Tangchi located in the northeast (Yuan et al., 2014). The lake is mainly replenished by groundwater (Zhu et al., 2016).

### 2.2 Sample Collection and Processing

Four different sampling points, YZN, YZM, YZS, and YZDH, were designed for this study. The coordinates of each sampling point are listed in Table 1. The sampling process started from the northern YZN and proceeded southwards. overlying water samples were collected at each sampling point and initially mixed in a water bucket to ensure homogeneity. The mixed water sample was then used to rinse a 10L white plastic container. After rinsing, the water sample was transferred into the white plastic container and stored in an insulated box with ice packs. Three parallel samples were collected at each sampling point. The physicochemical factors of the water samples were measured in the laboratory within 1 hour after collection.

**Table 1:** Sampling Point Coordinates

| Sampling Point | Center longitude | Center latitude. |
| --- | --- | --- |
| YZN | 103.01298204 | 24.94576633 |
| YZM | 103.00040769 | 24.89029096 |
| YZS | 102.99118104 | 24.86705819 |
| YZDH | 102.99359101 | 24.86032686 |

YZN: Yangzong lake north, YZM: Yangzong lake middle, YZS: Yangzong lake south, YZDH: Yangzong DAHE.

pH value, oxidation-reduction potential, and dissolved oxygen of water samples were measured in situ using a portable water quality analyzer (YSI 6600v2). Ammonia nitrogen was determined using the spectrophotometric method with Nessler’s reagent (HJ 535-2009), total phosphorus was measured using the spectrophotometric method with ammonium molybdate (GB/T11893-89), and total nitrogen was analyzed using the digestion-UV spectrophotometric method with alkaline potassium persulfate (HJ 636-2012). Arsenic content was measured using ICP-MS.

### 2.3 Microbial Sequencing Method for Water Samples

The samples for determining bacterial abundance and diversity were taken from the overlying water. There were four sampling points, and each sampling point had three replicates. High-throughput sequencing of 16S rDNA PCR products was carried out by Guangdong Gidio Biotechnology Co., Ltd. Genomic DNA was extracted from the samples, and the V3+V4 region of the 16S rDNA was amplified using specific primers with barcodes. The primer sequences used were 338F (5-ACTCCTACGGGAGGCAGCA-3) and 806R (5-GGACTACHVGGGTWTCTAAT-3). The amplification system consisted of a 50μL reaction mixture containing 5μL of 10× KOD Buffer, 5μL of 2.5mmol/L dNTPs, 1.5μL of primers (5μmol/L), 1 μL of KOD polymerase, and 100ng of template DNA. The amplification conditions were as follows: initial denaturation at 95°C for 2 minutes, followed by 27 cycles of denaturation at 98°C for 10 seconds, annealing at 62°C for 30 seconds, extension at 68°C for 30 seconds, and a final extension at 68°C for 10 minutes. The PCR amplification products were then gel extracted, and the concentration was measured using a QuantiFluorTM fluorometer. The purified amplification products were mixed in equal amounts and ligated with sequencing adapters, and the sequencing library was constructed according to the Illumina official protocol. The library was sequenced on the Hiseq2500 platform in PE250 mode to determine the relative abundance of the bacterial community in the overlying water.

### 2.4 Data Analysis

For data organization and preliminary analysis, Excel 2021 software (Microsoft Corporation, USA) was used. The data results were expressed as mean ± standard deviation (mean±SD). Spearman correlation analysis and significance tests were performed using SPSS 25.0 software (SPSS Inc., Chicago, IL, USA). Graphs were created using Origin 2019 software (OriginLab Corporation).

## 3 Results

### 3.1 Analysis of Differences in Physicochemical Properties of Water Samples

In this study, the pH of the overlying water ranged from highest to lowest as follows: YZN > YZS > YZDH > YZM. There were no significant differences in pH between the different sampling points (*P*>0.05). The highest levels of dissolved oxygen, total nitrogen, total phosphorus, and temperature were observed in the surface water of YZDH. YZM had the lowest levels of dissolved oxygen, temperature, and total phosphorus. YZN had the lowest total nitrogen content but the highest ammonia nitrogen content in the overlying water. YZM had the highest content of As, while YZN had the lowest. The order of As content from highest to lowest was YZM > YZDH > YZS > YZN. YZDH had the highest oxidation-reduction potential in the overlying water, and the oxidation-reduction potential at YZDH was significantly (P < 0.05) higher than that at YZM, YZS, and YZN. YZN had the lowest oxidation-reduction potential in the overlying water. Table 2 for specific details.

**Table 2:** Physical and Chemical Properties of Samples

| Points | pH | ORP | DO | T | NH<sub>3</sub>-N | TP | TN | As |
| --- | --- | --- | --- | --- | --- | --- | --- | --- |
| YZN | 8.275 ± 0.26a | -83.06 ± 3.25c | 0.86 ± 0.02a | 18.06 ± 1.84a | 0.58 ± 0.29a | 0.04 ± 0.02a | 0.84 ± 0.40a | 23.44 ± 6.65a |
| YZM | 8.075 ± 0.24a | 137.8 ± 3.91b | 0.85 ± 0.06a | 17.60 ± 1.58a | 0.44 ± 0.18a | 0.03 ± 0.01a | 0.87 ± 0.38a | 27.34 ± 8.16a |
| YZS | 8.093 ± 0.07a | 112.4 ± 35.24b | 0.87 ± 0.11a | 18.20 ± 1.60a | 0.48 ± 0.14a | 0.04 ± 0.01a | 0.97 ± 0.35a | 25.06 ± 7.24a |
| YZDH | 8.093 ± 0.15a | 199 ± 5.86a | 0.91 ± 0.12a | 18.96 ± 1.41a | 0.34 ± 0.06a | 0.05 ± 0.01a | 0.98 ± 0.27a | 26.05 ± 7.06a |

DAHE, ORP: Oxidation-Reduction Potential, DO: Dissolved oxygen, T: Temperature, AS: Arsenic, TP:
Total phosphorus, NH<sub>3</sub>-N: Ammonia nitrogen, TN: Total Nitrogen. a、b、c and d indicate significant differences.

### 3.2 The Composition of Bacterial Community Structure in the Overlying Water of Yangzong Lake

In the water samples, a total of 1,937,340 valid sequences were obtained, with an average of 80,722 valid sequences per sample. The total number of OTUs detected in the samples was 18,352. The Venn diagram of species showed that there were 743 shared OTUs among the sampling points(Figure 1). Among them, the northern sediment of Yangzong Lake had 593 unique OTUs, the central sediment of Yangzong Lake had 1,575 unique OTUs, the southern sediment of Yangzong Lake had 792 unique OTUs, and the sediment of the Yangzong DAHE had 1,064 unique OTUs.

**Figure 1:**
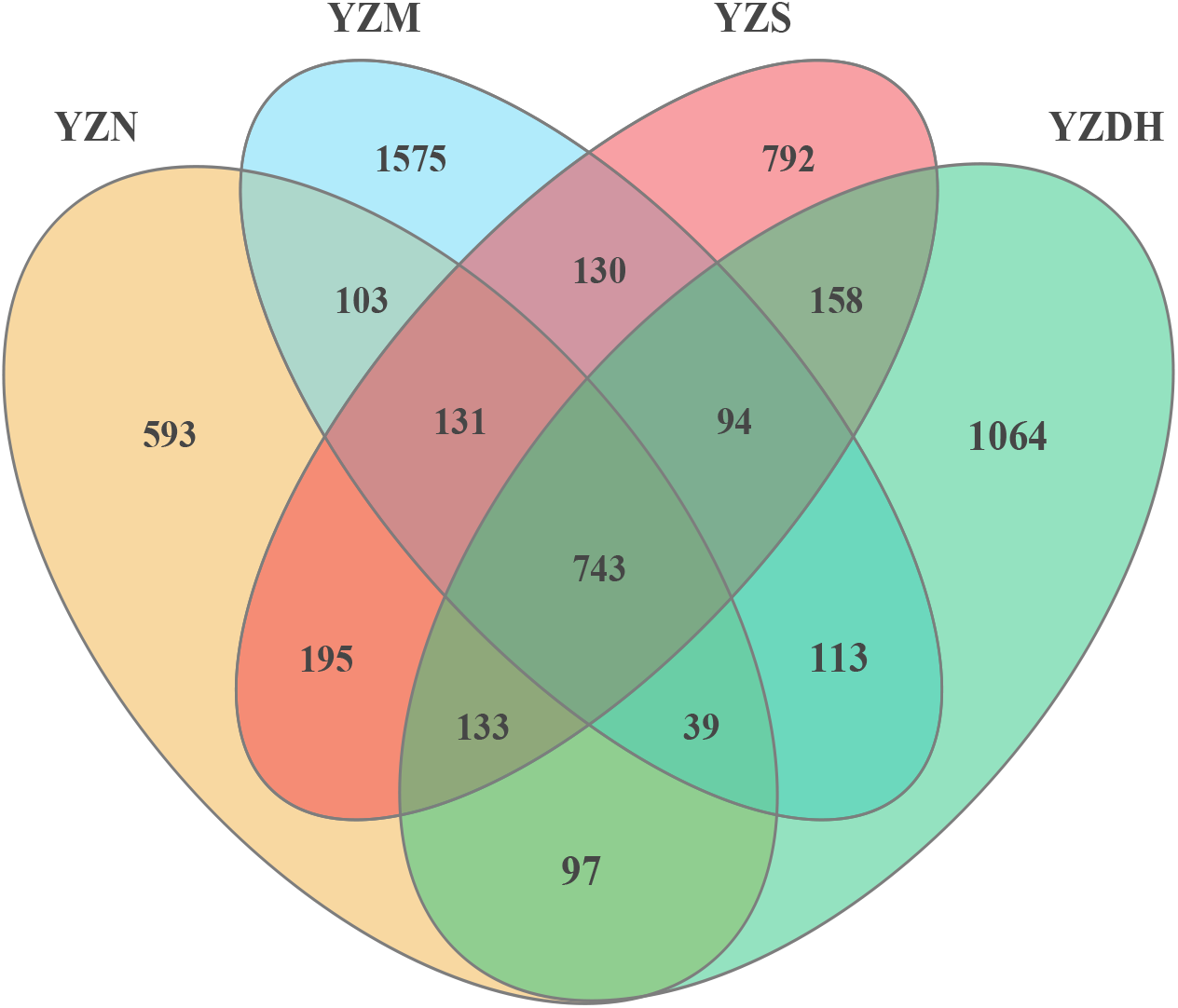
Venn diagram of species distribution.YZN: Yangzong lake north, YZM: Yangzong lake middle, YZS: Yangzong lake south, YZDH: Yangzong DAHE

The results were presented using the Sob, Chao, Shannon, and Simpson indices in Table 3. The Sob index represents the number of OTUs obtained during the sequencing process, with higher values indicating greater species diversity in the sample. The species diversity in the overlying water of Yangzong Lake was significantly higher (P<0.05) than in the overlying water of the southern part of the lake.

**Table 3:**
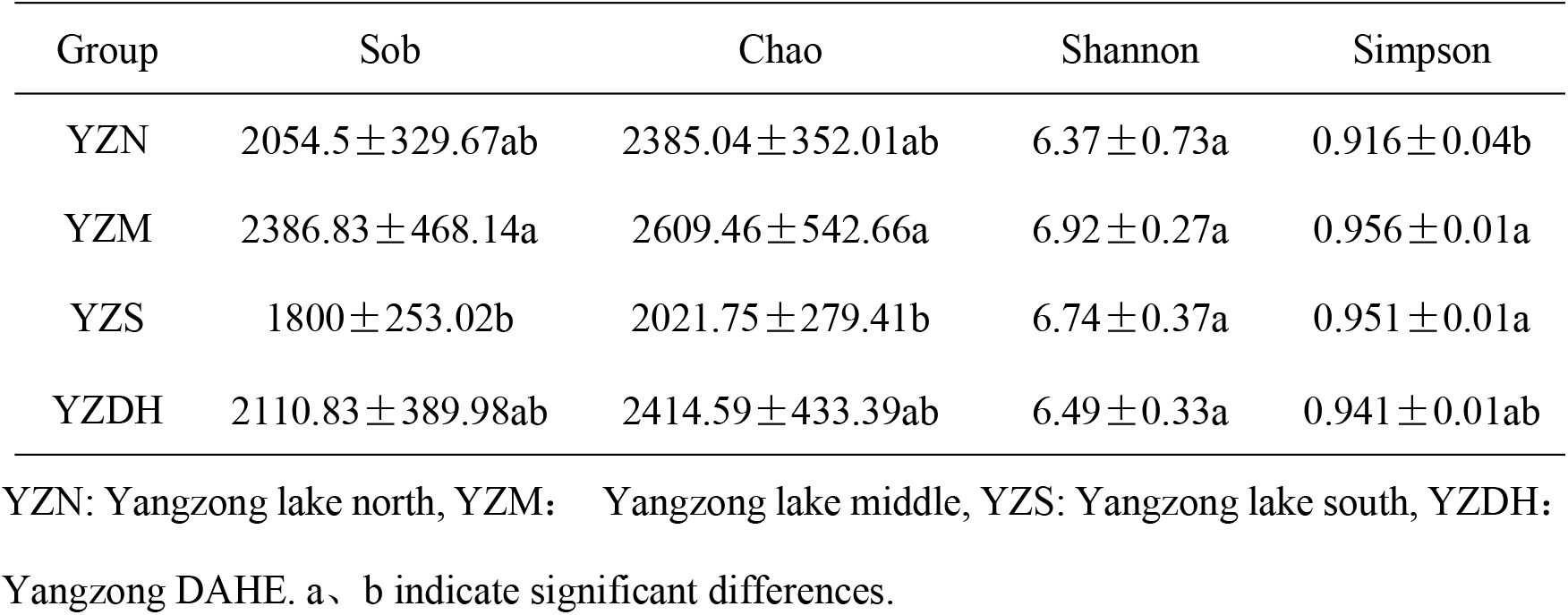
Analysis of bacterial alpha diversity indices.

The Chao index reflects the richness of the bacterial community, with higher values indicating greater abundance of bacterial species. The richness of the bacterial community in the overlying water of Yangzong Lake was significantly higher (P<0.05) than in the northern part of the lake and the Yangzong DAHE, and it was also higher than in the bacterial community of the southern part of Yangzong Lake.

The Simpson and Shannon indices integrate the abundance and evenness of species. In this study, the Simpson and Shannon indices ranked from high to low as follows: YZM > YZS > YZDH > YZN. However, the differences between the samples were not significant.

### 3.3 Analysis of bacterial community structure in the overlying water of Yangzonghai Lake

The top ten abundant bacterial communities at the phylum level in the overlying water of Yangzong Lake are Actinobacteriota, Bacteroidota, Proteobacteria, Cyanobacteria, Verrucomicrobiota, Patescibacteria, Chloroflexi, Campylobacterota, Planctomycetota, and Firmicutes (refer to Figure 2 for specific details).

**Figure 2:**
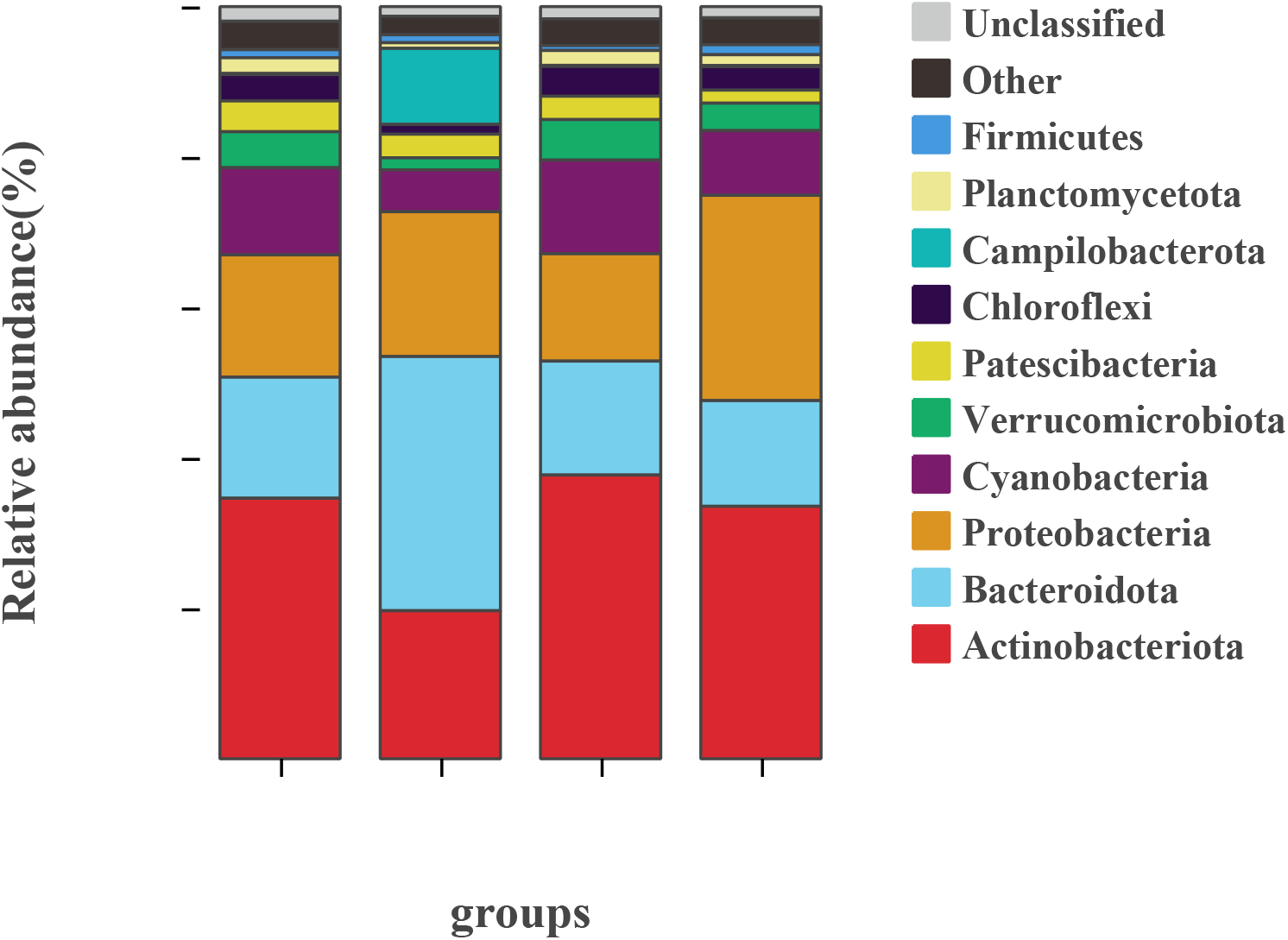
Distribution of bacterial phyla at the genus level in the overlying water. YZN: Yangzong lake north, YZM: Yangzong lake middle, YZS: Yangzong lake south, YZDH: Yangzong DAHE

Actinobacteriota was the most abundant phylum, with relative abundances ranging from 33.56% to 37.71% In YZN?YZS?YZDH. In the overlying water of the YZM, Bacteroidota was the most abundant phylum, with a relative abundance of 33.79%.

The top ten abundant bacterial genera at the species level in the overlying water (refer to Figure 3 for specific details) are hgcI_clade, CL500-29_marine_group, Flavobacterium, Acinetobacter, Terrimonas, Cylindrospermopsis_CRJ1, Sediminibacterium, Prochlorothrix_PCC-9006, Limnohabitans, and Pseudarcicella.

**Figure 3:**
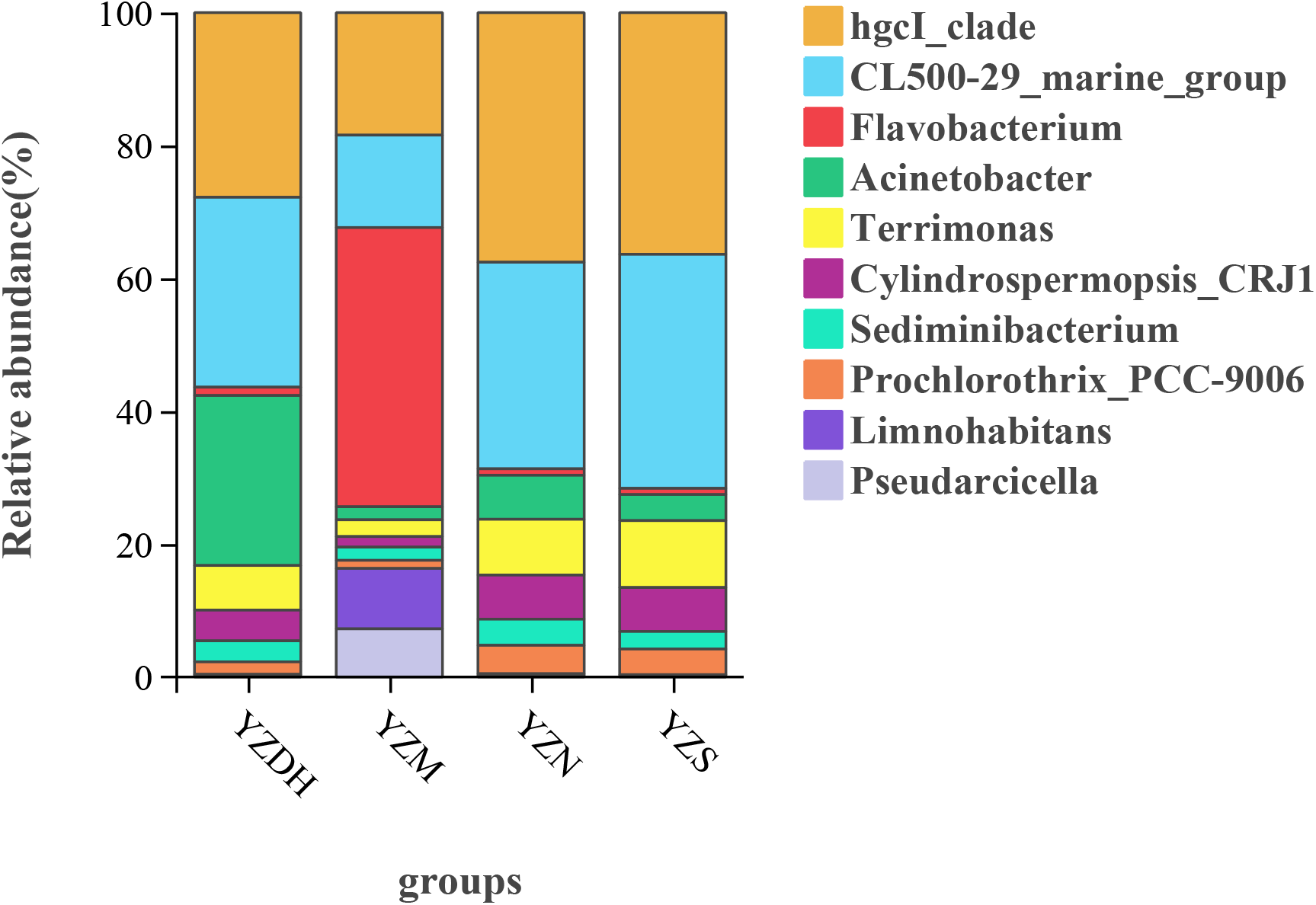
Distribution of bacterial genera at the species level in the overlying water. YZN: Yangzong lake north, YZM: Yangzong lake middle, YZS: Yangzong lake south, YZDH: Yangzong DAHE

The genus hgcI_clade had the highest relative abundance, reaching 37.56% In YZN YZS. In the overlying water of YZM, the genus Flavobacterium had the highest relative abundance, reaching 42.02%. In the overlying water of YZDH, the genus CL500-29_marine_group had the highest relative abundance, reaching 28.57%.

Based on β diversity analysis, it was found that there is a significant difference in the bacterial community structure of the overlying water in YZM compared to other sampling sites ( Figure 4). ANOSIM and ADONIS tests were performed to assess the inter-group differences, and the results showed a significant difference in the bacterial community structure among the sampling points at a significance level of (P<0.05). This indicates that there are significant differences in the composition and relative abundance of bacterial communities among the different sampling points.

**Figure 4:**
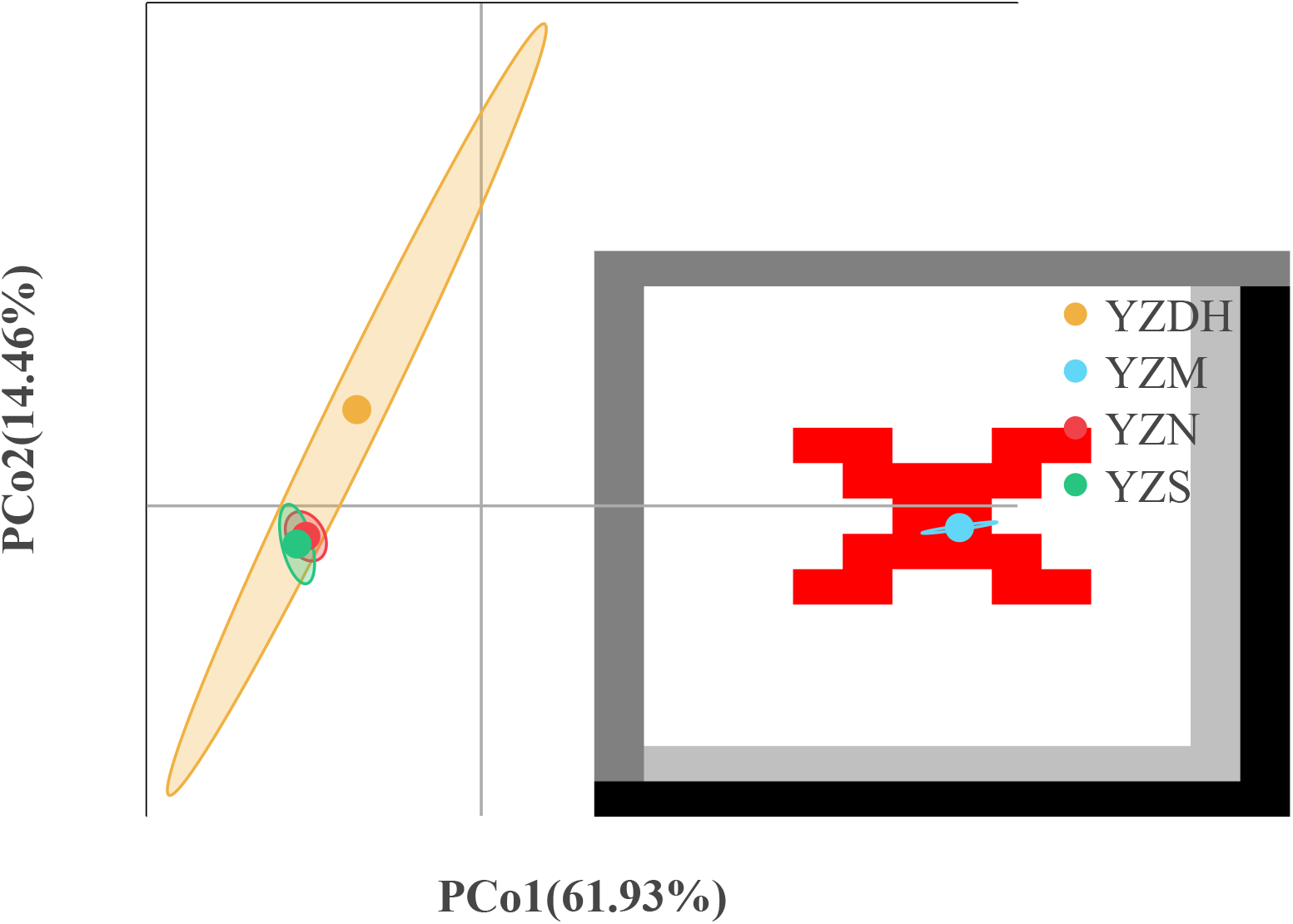
Principal Component Analysis (PCA) plot of the bacterial community structure. YZN: Yangzong lake north, YZM : Yangzong lake middle, YZS: Yangzong lake south, YZDH : Yangzong DAHE.

### 3.4 Correlation Analysis Between Water Environmental Factors and Bacterial Diversity

pH, ORP, DO, temperature, and AS were positively correlated with Sob, Chao, Shannon, and Simpson indices ( Figure 5). Specifically, AS had a significant positive correlation with the Sob index. On the other hand, TP was negatively correlated with the Sob, Chao, Shannon, and Simpson indices. NH_3_-N and TN were negatively correlated with the Sob and Chao indices but positively correlated with the Shannon and Simpson indices.

**Figure 5:**
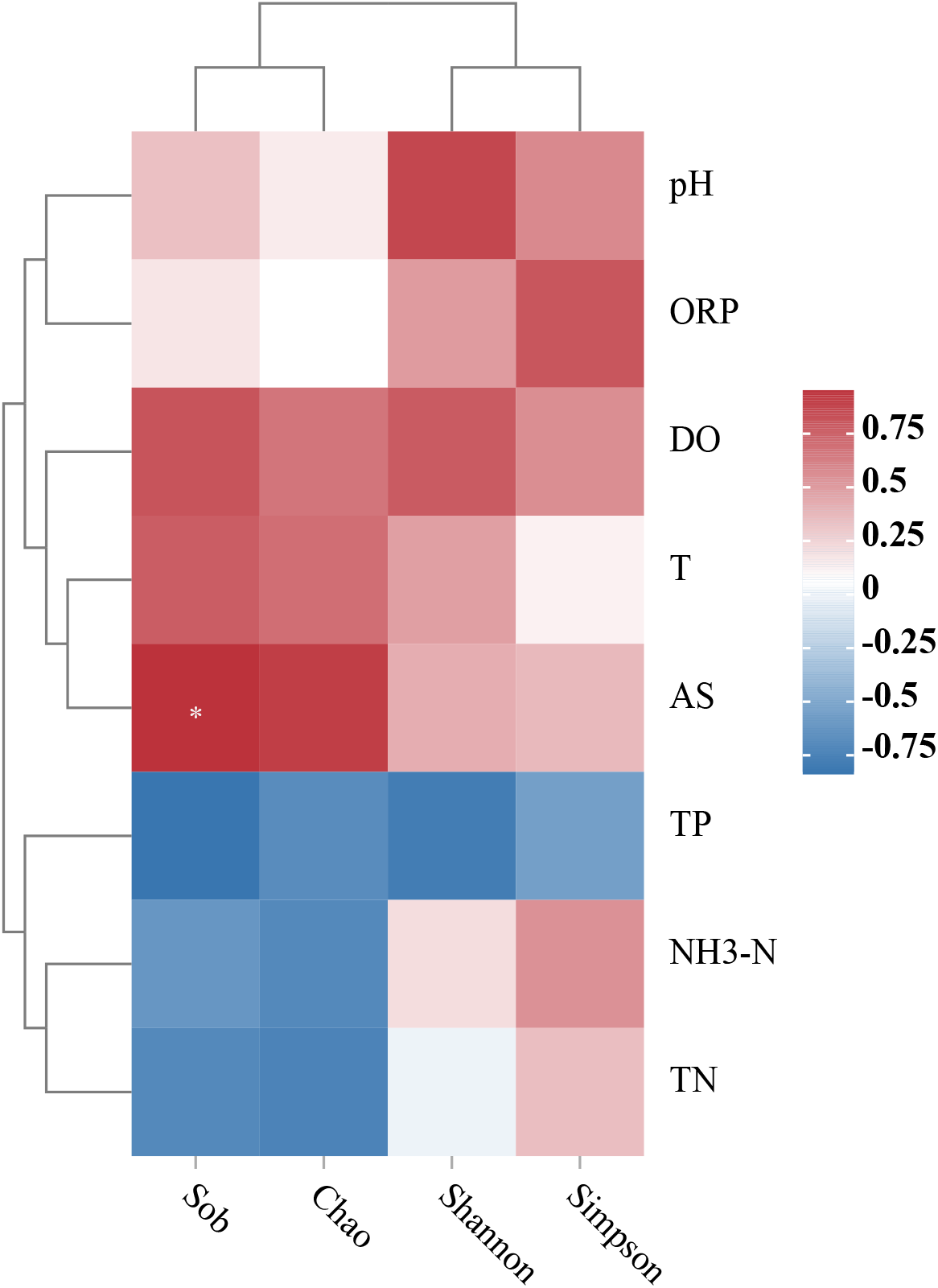
Correlation analysis between environmental factors and bacterial α diversity.ORP : Oxidation-Reduction Potential, DO: Dissolved oxygen, T: Temperature, AS: Arsenic, TP: Total phosphorus, NH_3_-N: Ammonia nitrogen, TN: Total Nitrogen.

### 3.5 Correlation Analysis between Water Environmental Factors and Bacterial Community Structure

This study analyzed the correlation between bacterial phylum and genus levels and environmental factors. At the phylum level ( Figure 6), Actinobacteriota and Chloroflexi showed a positive correlation with NH_3_-N, TP, and TN, while they showed a negative correlation with pH, ORP, DO, temperature (T), and AS. Bacteroidota and Campilobacterota showed a positive correlation with pH, ORP, DO, T, and AS, while they showed a negative correlation with NH_3_-N, TP, and TN. Additionally, Campilobacterota showed a highly significant positive correlation with DO (p<0.001) and a highly significant negative correlation with TP (p<0.0011. Proteobacteria and Firmicutes showed a positive correlation with pH and T, while they showed a negative correlation with ORP, NH_3_-N, TN, and AS. Cyanobacteria, Verrucomicrobiota, and Planctomycetota showed a positive correlation with NH_3_-N, TP, TN, and ORP, while they showed a negative correlation with pH, DO, T, and AS. Patescibacteria showed a positive correlation with pH, ORP, DO, NH_3_-N, and TN, while they showed a negative correlation with TP, T, and AS.

**Figure 6:**
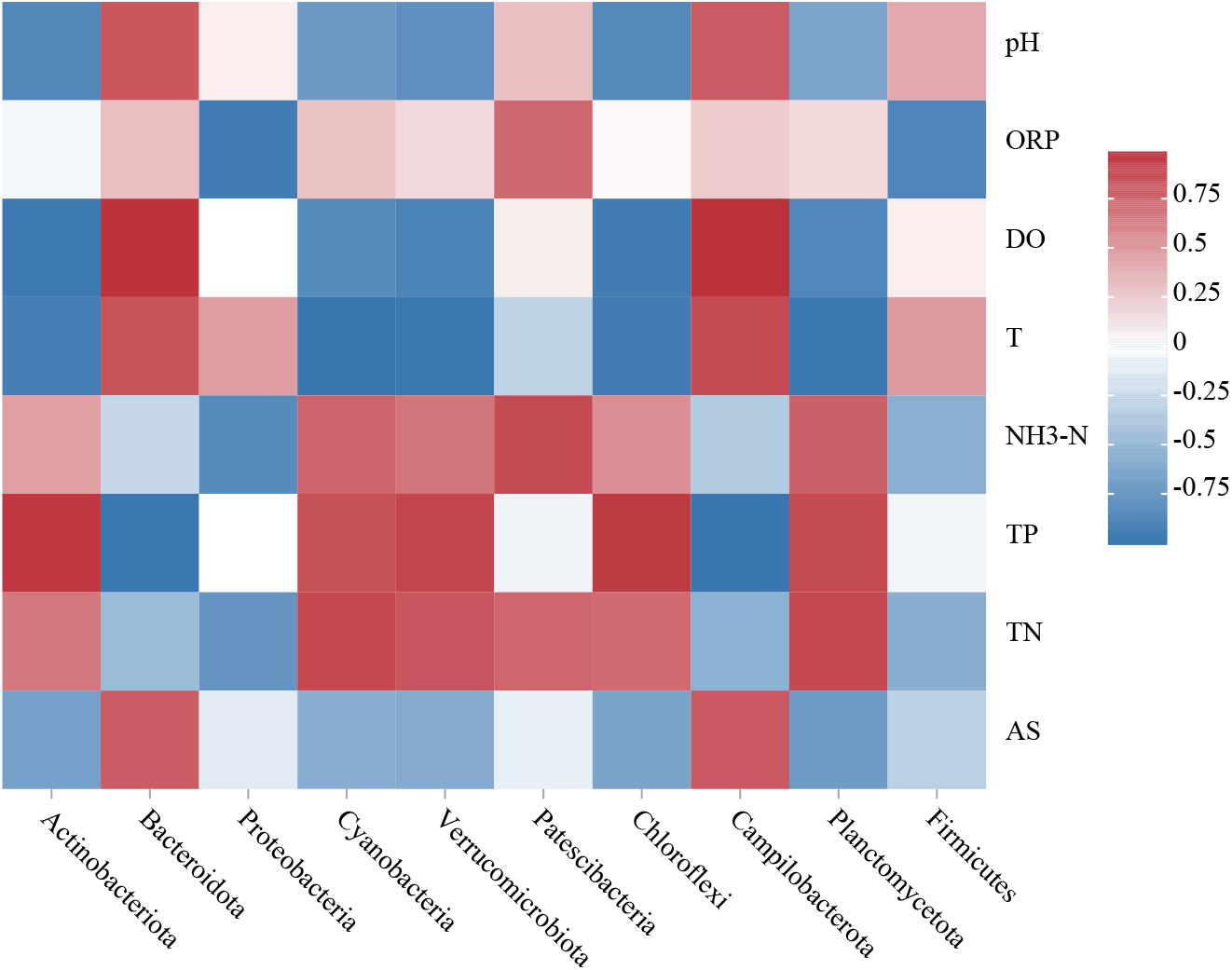
Analysis of the correlation between water environmental factors and bacterial phylum-level community structure. ORP: Oxidation-Reduction Potential, DO: Dissolved oxygen, T: Temperature, AS: Arsenic, TP: Total phosphorus, NH_3_-N: Ammonia nitrogen, TN: Total Nitrogen

At the genus level (Figure 7), hgcI_clade, Cylindrospermopsis_CRJ1, CL500-29_marine_group, Terrimonas, and Prochlorothrix_PCC-9006 showed a negative correlation with pH, temperature (T), dissolved oxygen (DO), and AS, while they showed a positive correlation with ORP, NH_3_-N, TN, and TP. Sediminibacterium showed a negative correlation with pH, T, DO, AS, and ORP, while it showed a positive correlation with NH_3_-N, TN, and TP. Acinetobacter showed a negative correlation with pH, T, DO, AS, ORP, NH_3_-N, and TN, while it showed a positive correlation with TP. Flavobacterium, Limnohabitans, and Pseudarcicella showed a positive correlation with pH, T, DO, AS, and ORP, while they showed a negative correlation with NH_3_-N, TN, and TP. Among them, Flavobacterium, Limnohabitans, and Pseudarcicella exhibited a highly significant positive correlation with DO (p<0.001) and a highly significant negative correlation with TP(p<0.001).

**Figure 7:**
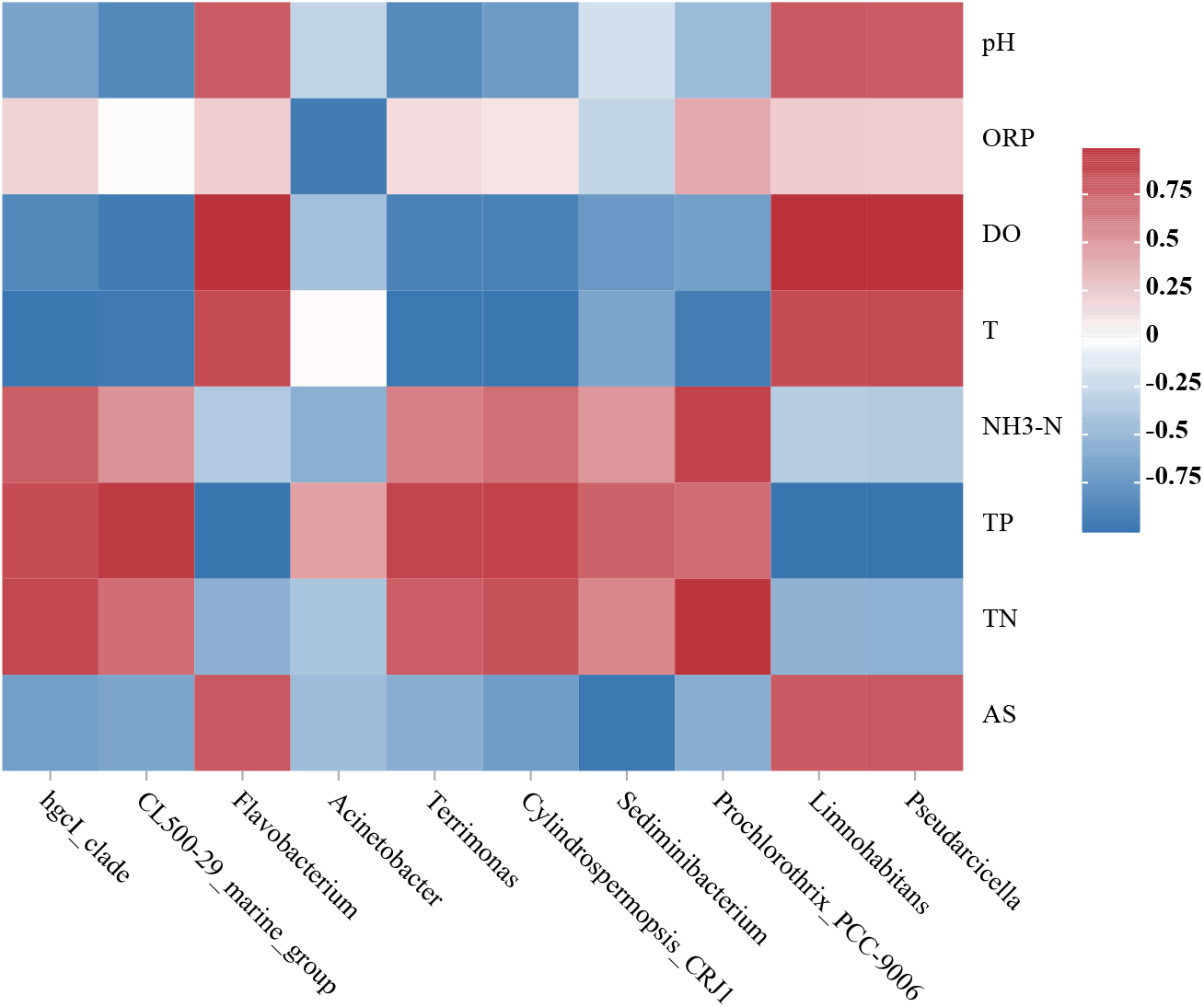
Analysis of the correlation between water environmental factors and bacterial genus-level community structure. ORP : Oxidation-Reduction Potential, DO : Dissolved oxygen, T : Temperature, AS: Arsenic, TP: Total phosphorus, NH_3_-N: Ammonia nitrogen, TN: Total Nitrogen.

## 4 Discussion

### 4.1 Analysis of Differences in Physicochemical Factors of the Overlying Water in Yangzong Lake

In this study, an analysis was conducted on the physicochemical factors of the overlying water in YZN and YZM, focusing on the redox potential (ORP). The ORP is an indicator reflecting the redox status of the water or soil (Li et al., 2004; Gao et al.,2002). An ORP value greater than 0mV indicates an oxidative medium, while a value less than 0mV indicates a reducing medium (Khumalo et al., 2022).

In this study, it was found that the ORP of the overlying water in YZN was less than 0mV, indicating its reducing nature. This is attributed to the release of iron in the sediment of Yangzong Lake, leading to a decrease in ORP due to an increase in iron release (Jing et al., 2014). Additionally, the magnitude of ORP is influenced by factors such as water temperature, pH, dissolved oxygen, etc. (Ma et al., 2020). An increase in pH leads to a decrease in ORP value (Köse et al., 2014), and an increase in temperature also causes a decrease in ORP value (Yin et al., 2005). The thermal discharge from coal-fired power plants on the northern coast of Yangzong Lake may also be a contributing factor to the variation in ORP (Zhai et al., 2010).

Furthermore, the overlying water in YZM exhibited the lowest dissolved oxygen content and water temperature. This can be attributed to the greater depth of Yangzong Lake at the YZM location, which aligns with the conclusions drawn from studies on the vertical physicochemical factors of water bodies in Fuxian Lake and Zurich Lake (Kim et al., 2006; Zhou et al., 2022).

### 4.2 Analysis of bacterial diversity and community structure in the overlying water of Yangzong Lake

The α-diversity of bacteria is an indicator reflecting the abundance and community structure of bacterial populations. In this study, the diversity index of the overlying water in YZM was significantly higher than that of other sampling sites, indicating a greater species richness and more complex community structure of bacteria in YZM (Wang et al., 2023).

Bacterial α-diversity can be influenced by human activities. Studies by Wang et al. have shown that human activities can reduce the diversity index of bacteria, and the development of tourism can lower the α-diversity index of bacteria in wetland parks (Wang et al., 2020; Wang et al., 2020 ). In this study, YZM is located in the central part of Yangzong Lake and experiences less human disturbance compared to other sampling sites. Therefore, its diversity index is significantly higher than that of other sites.

The bacterial communities with a relatively high abundance at the phylum level in the overlying water of Yangzong Lake include Actinobacteriota, Bacteroidota, Proteobacteria, and Cyanobacteria. These phyla are also predominant in arsenic-contaminated water (Davolos et al., 2011), which is consistent with other findings. (Zhang et al., 2019). Actinobacteriota has been found to have a higher relative abundance in different water layers of deep lakes (Okazaki et al., 2016). The higher relative abundance of Actinobacteriota in the overlying water of Yangzong Lake may be attributed to their smaller cell size and higher utilization efficiency of low nutrient concentrations (Salcher et al., 2011). Proteobacteria encompass a wide range of autotrophic and heterotrophic bacteria that are commonly found in aquatic environments. Proteobacteria and Bacteroidota are abundant bacterial groups in lake environments, which is consistent with other research findings (Gao et al., 2022).Cyanobacteria are capable of both photosynthesis and nitrogen fixation and have been widely observed in reservoirs and lakes, similar to the results of this study (Sanseverino et al., 2022; Li et al., 2022).The genera HgcI_clade, CL500-29_marine_group, and Flavobacterium exhibit relatively high abundance at the genus level in the overlying water of Yangzong Lake. HgcI_clade belongs to the phylum Actinobacteriota, class Actinobacteria, order Frankiales, and family Sporichthyaceae. It plays a role in promoting the cycling of carbon and nitrogen in water environments, improving water quality. HgcI_clade is considered one of the persistent microorganisms in global oligotrophic aquatic ecosystems, characterized by its high and stable abundance (Huang et al., 2020; Li et al., 2016). Previous studies have shown that HgcI_clade is a dominant bacterial group in water environments of rice paddies under different fertilization modes and fish-vegetable co-culture systems (Newton et al., 2011; Ruprecht et al., 2021), which is consistent with the results of this study. CL500-29_marine_group belongs to the phylum Actinobacteriota, class Acidimicrobiia, order Microtrichales, and family Ilumatobacteraceae. It plays a dominant role in the evolution of water quality (Zhu et al., 2020; Chen et al., 2017). Both CL500-29_marine_group and HgcI_clade, belonging to the phylum Actinobacteriota, are common dominant genera in freshwater ecosystems. Moreover, they show similar spatiotemporal variations (Ruprecht et al., 2019; Liu et al., 2020).Flavobacterium belongs to the phylum Bacteroidota, class Bacteroidia, order Flavobacteriales, and family Flavobacteriaceae. It is known as a denitrifying bacterium that can degrade various high-molecular-weight compounds such as proteins, carbohydrates, and lipids, thereby removing nitrogen and phosphorus from the water (Zhou et al., 2020). This result is consistent with similar studies conducted in Jiangsu’s Lake Tai and Hubei’s Lake Liangzi (Chang et al., 2020; Wang et al., 2022).

### 4.3 Correlation analysis of environmental factors and bacterial community structure Actinobacteriota and Chloroflexi are positively correlated with NH_3_-N, TP, and TN because

phosphorus (P) and nitrogen (N) are essential nutrients and energy sources for bacterial growth and metabolism. The presence of NH_3_-N, TP, and TN is a prerequisite for bacterial growth and metabolism (Zhang et al., 2020). In this study, Actinobacteriota and Chloroflexi are negatively correlated with AS (heavy metals) because heavy metals induce the generation of reactive oxygen species, which can be toxic to bacteria (Li et al., 2017).

The environmental factors pH, ORP, water temperature, dissolved oxygen (DO), total nitrogen (TN), total phosphorus (TP), and heavy metal content are driving factors influencing the differences in bacterial community structure in lake ecosystems (Wang et al., 2023; Roguet et al., 2015; Zhong et al., 2016).

In this study, Flavobacterium, Limnohabitans, and Pseudarcicella were found to have a highly significant correlation (p<0.001) with DO and TP. TP is considered a crucial determinant of bacterial community structure in lake water and sediments (Chen et al., 2023). Flavobacterium, Limnohabitans, and Pseudarcicella, belonging to the phylum Bacteroidetes and class Betaproteobacteria, were identified as dominant species and were significantly influenced by TP, which is consistent with the study’s findings (Wang et al., 2021). These bacteria, including Flavobacterium, Limnohabitans, and Pseudarcicella, are involved in the decomposition and cycling of substances in the water environment (Zheng et al., 2022; Wu et al., 2019). As a result of their involvement in decomposition and cycling processes, they consume oxygen, which may explain the strong correlation with DO.

The PCoA analysis revealed significant differences in the bacterial community composition between the YZM point and other sampling sites. These differences might be attributed to the environmental factors and hydrological conditions of the lake. For instance, YZM exhibited significantly lower dissolved oxygen levels and water temperature compared to other sites. Dissolved oxygen and water temperature are known to be key factors driving the variations in bacterial community composition in lakes (Wang et al., 2020; He et al., 2021).

## 5 Conclusion

1. There are significant differences in the oxidation-reduction potential (ORP) of the overlying water at different sampling points in Yangzong Lake. The other physicochemical indicators of water quality show noticeable differences but do not exhibit significance.
2. The number of OTUs and bacterial alpha diversity index in the surface water of YZM are significantly higher than at other sampling points.
3. The dominant bacterial phyla in the overlying water of Yangzong Lake are Actinobacteriota, Proteobacteria, and Bacteroidota. The dominant bacterial genera include hgcI_clade, CL500-29_marine_group, and Flavobacterium. Dissolved oxygen (DO) and total phosphorus (TP) are identified as the major environmental factors influencing bacterial community structure.

